# The structure of the SufBC_2_D-SufE complex reveals the mechanism of sulfur transfer in bacterial Fe-S cluster assembly

**DOI:** 10.64898/2026.05.18.725997

**Authors:** Nidhi Chhikara, Nadia Mireku, Jack A. Dunkle, Patrick A. Frantom

**Author notes:** to whom correspondence should be addressed: Patrick A. Frantom,; Jack A. Dunkle.

## Abstract

Iron-sulfur clusters are essential cofactors assembled in bacteria by the Suf pathway through a series of transient protein-protein interactions that transfer sulfur from L-cysteine to a scaffold complex. While early steps in persulfide transfer are well characterized, the mechanism of sulfur delivery to the SufBC_2_D scaffold has remained unresolved. Here, we report the first structure of the SufBC_2_D-SufE complex, capturing the final step in persulfide transfer in the Suf pathway. The structure reveals coordinated conformational changes in both SufB and SufE that expose the otherwise buried C254 acceptor site and position the SufE C51 loop beneath the SufB-SufD axis. Biochemical analysis of SufB variants demonstrates that substitutions in the globally conserved 220s β-strand enhance SufE binding affinity and persulfide transfer rates, consistent with stabilization of a locally rearranged, transfer-competent conformation. Together, these results support a model in which conformational gating regulates persulfide transfer, providing a mechanism for controlling access to reactive sulfur intermediates.

## Introduction

Iron-sulfur (Fe-S) clusters are essential metallocofactors required for core biological processes including cellular respiration, DNA replication, and signaling.(1–3) Although Fe-S clusters can often be assembled spontaneously in vitro on apo-target proteins by addition of iron, sulfide, and a reductant, in vivo cluster biogenesis in most organisms depends on dedicated multi-protein pathways that regulate cluster assembly and trafficking.(4) Common features of these pathways include a PLP-dependent cysteine desulfurase and a central cluster assembly scaffold. The cysteine desulfurase cleaves the C-S bond of L-cysteine to generate L-alanine and a covalent persulfide intermediate on an active site cysteine residue (R-S-SH).(5, 6) This persulfide must then be transferred to the assembly scaffold, which also coordinates acquisition of iron and reducing equivalents to promote formation of nascent Fe-S clusters. As the process of Fe-S cluster assembly is essential in bacteria, including major human pathogens(7, 8), these pathways represent attractive targets for the development of new antibiotic therapeutics.

Bacteria have evolved several Fe-S cluster assembly pathways, including major housekeeping systems such as the iron sulfur cluster (Isc) and the sulfur formation (Suf) pathways, the specialized nitrogen fixation (Nif) pathway for maturation of nitrogenase, and the recently identified minimal systems for Isc (Mis) and Suf (Sms) pathways, which may represent early evolutionary forerunners of their cognate systems.(9–12) The best characterized pathway for bacterial Fe-S cluster assembly is the Isc pathway, likely due to its central role in *Escherichia coli* metabolism and its evolutionary relationship to the mitochondrial ISC pathway. Here, the cysteine desulfurase IscS and the scaffold protein IscU form a complex that allows for direct transfer of the cysteine-derived persulfide, located on a long, flexible loop, from the desulfurase to the assembly scaffold in a process stimulated by iron occupancy on IscU (Figure 1A).(13)

**Figure 1.**
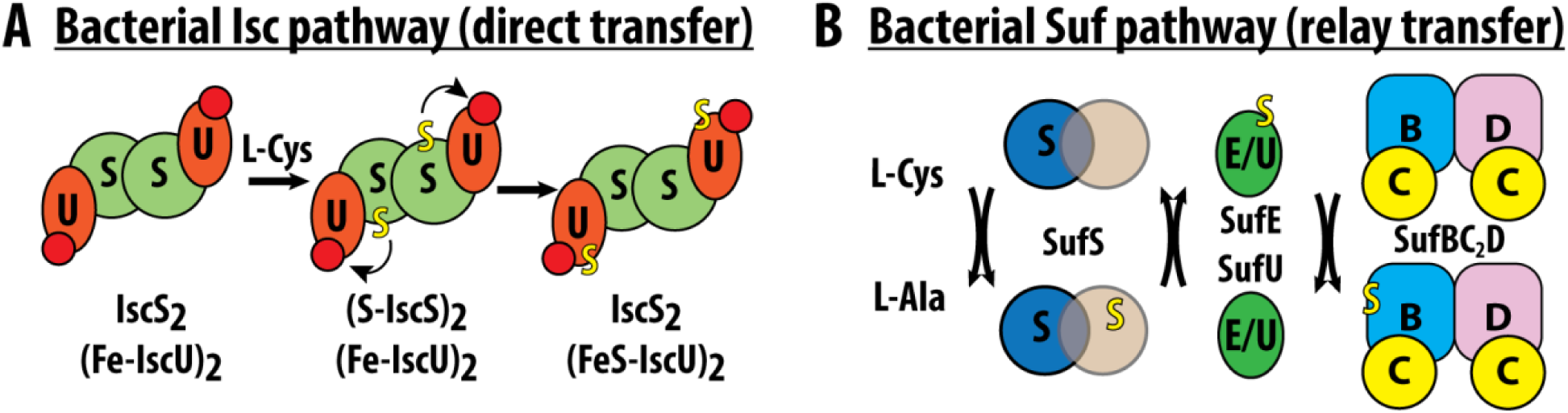
Bacterial Fe-S cluster assembly pathways. (A) Persulfide transfer in the bacterial Isc pathway is direct from IscS to IscU. This step is stimulated by iron (red circle) occupancy on IscU. (B) Persulfide trafficking in the bacterial Suf pathway occurs via multiple transfer steps between cysteine residues on SufS, SufE or SufU, and SufBC_2_D.

Although the Isc pathway is the best understood, the Suf pathway is most prevalent in bacteria.(14) The Suf pathway assembles clusters using SufS as a cysteine desulfurase and the heterotetrameric SufBC_2_D complex as an assembly scaffold.(15, 16) In contrast to the Isc pathway, there is no evidence for direct interaction between SufS and SufBC_2_D for persulfide transfer. Instead, an additional transpersulfurase protein (SufE or SufU) is required to accept the persulfide from SufS and relay it to the SufB subunit of the SufBC_2_D complex (Figure 1B). This distinctive pathway architecture, in which a covalent intermediate is passed sequentially among three proteins without formation of a stable ternary complex or covalent tether, implies the existence of stringent regulatory mechanisms to prevent formation of dead-end complexes and enforce directional sulfur transfer from desulfurase to assembly scaffold.

The first stage of persulfide trafficking in the Suf system has been well described, and SufS enzymes from both SufE- and SufU-dependent pathways share substantial structural and mechanistic homology. In Gram-positive systems, like *Bacillus subtilis*(*17*), *Mycobacterium tuberculosis*(*18*) and *Staphylococcus aureus*(*19*), SufU functions as the transpersulfurase and requires a zinc ion for activity. The zinc coordinates the SufU active site cysteine that accepts the persulfide, and the zinc coordination site undergoes a ligand swap with a histidine residue on SufS to regulate persulfide transfer in these systems.(20) In *E. coli*, and other Gram-negative organisms, SufU is replaced by the metal-independent SufE. Persulfide transfer from SufS to SufE requires conformational changes on SufS linked to specific catalytic intermediates in the desulfurase reaction.(21–23) These conformational changes promote formation of a “close approach” complex between SufS and SufE that enables persulfide transfer. Both transpersulfurases utilize a strictly conserved arginine residue (R125 in *B. subtilis* SufU, R119 in *E. coli* SufE) that senses completion of persulfide transfer and contributes to dissociation of the SufS-transpersulfurase complex.(22, 24) These observations support a general model in which conformational gating, coupled to catalytic intermediate formation, regulates the trafficking of reactive persulfide intermediates by controlling transient protein-protein interactions and enforcing directional sulfur flux.

The second stage of persulfide trafficking, from the transpersulfurase to the SufBC_2_D assembly scaffold, remains poorly understood. The only system characterized mechanistically for this step is the *E. coli* Suf pathway, which is SufE-dependent and utilized as an emergency system induced during times of oxidative stress or iron starvation.(25) Biochemical studies support a model in which persulfide transfer occurs from C51 on SufE to C254 on the SufB subunit of SufBC_2_D, as sulfur transfer and persulfide accumulation are diminished in C254A SufB variants.(26–28) Accordingly, C254 is generally accepted to be the initial persulfide acceptor site on the scaffold complex. Despite this biochemical framework, the structural mechanism for this step remains unclear. To date, only two X-ray crystal structures of the complex have been reported, including a 4.28 Å structure and a 2.96 Å mercury-derivatized structure.(26) In both structures, substantial regions of SufB are unresolved, including a flexible segment adjacent to the C254-containing pocket (residues 78-156). Although the missing density creates the appearance of an accessible opening toward C254, the unresolved region likely contributes to regulating access to this pocket in the intact complex, leaving unclear how the transpersulfurase gains productive access for sulfur transfer (Figure S1).

Several experimental observations further suggest that the mechanism is more complex than initial transfer to C254 alone. In vivo persulfide accumulation on SufBC_2_D can be decreased not only by mutation of C254, but also by mutation of another conserved cysteine, C405, located at the SufB-SufD interface.(29) C405 is positioned near the proposed site of Fe-S cluster assembly, and previous structural analyses identified an internal tunnel connecting C254 and C405 in SufB.(26) However, there is currently no clear mechanism by which a covalent persulfide species could traverse this tunnel in the absence of intervening cysteine residues or a defined reductive step to generate S^2-^.

In this report, we provide direct structural evidence that C254 is the initial site of persulfide acceptance from SufE and identify local conformational rearrangements that promote persulfide transfer. Using sequence conservation analysis and biochemical assays, we show that alanine substitutions at conserved residues Y224, R226, or N228 on SufB produce SufBC_2_D complexes with accelerated rates of persulfide acceptance from SufE relative to the wild-type complex, together with an approximately 7-8-fold increase in affinity for SufE, consistent with stabilization of a locally rearranged, transfer-competent conformation of SufB. Leveraging this enhanced interaction, single particle cryo-electron microscopy was used to capture a structure of the Y224A SufBC_2_D variant in a complex with SufE, which was refined to 4.2 Å resolution. Analysis of the resulting structural model identifies conformational changes in both SufB and SufE that position C51 and C254 within 8 Å of one another in an extended configuration compatible with persulfide transfer. These results define the initial step of sulfur delivery from SufE to the scaffold and support a model in which conformational gating of SufB and SufE regulates productive persulfide transfer, thereby enforcing directional sulfur trafficking in the Suf pathway.

## Results

### Mapping of conserved surface residues on SufB

We hypothesized that the SufB interface for persulfide transfer would be strongly conserved throughout evolution. To generate an unbiased multiple-sequence alignment that captured the sequence diversity of the SufB fold, the Enzyme Function Initiative-Enzyme Similarity Tool (EFI-EST) was used to create a sequence similarity network for visualization in Cytoscape. Application of an edge alignment score cutoff of 220 resulted in a final network containing 1,642 nodes connected by 20,258 edges (Figure S2A). The resulting clusters are largely separated around taxonomy, including clear distinctions between Bacillati and Pseudomonadati, likely reflecting use of SufU or SufE transpersulfurases, respectively. Eukaryotic sequence clusters were still connected to the major Pseudomonadati cluster, consistent with SufE usage in eukaryotic plastid organelles.(30) Sequences for Archaea also separated into unique clusters.

Representative sequences from major clusters and sub-clusters in the network were selected to generate an unbiased 23-sequence multiple sequence alignment using MAFFT (Figure S2B). Sequence identity at each position was mapped onto the SufB subunit of the SufBC_2_D complex (Figure 2A). Among a handful of conserved exterior regions, a patch of globally conserved residues (224–228) was identified on the 220s beta strand directly exterior to C254 (Figure 2B and 2C). Three of these residues have previously been investigated with R226A and N228A substitutions leading to loss of complementation growth phenotypes, while substitution of Y224A resulted in a partial, temperature sensitive complementation.(29) These phenotypes support an important functional role for these globally conserved residues in Fe-S cluster assembly. *Kinetic characterization of persulfide transfer from SufS to SufBC_2_D variants.* To better determine functional roles for SufB residues 224-228, site-directed mutagenesis was used to introduce alanine substitutions in a previously described pDuet vector for co-expression of the *sufBCD* genes from a single promoter with an *N*-terminal (His)_6_-tag on SufB.(31) Expression from this construct results in purification of SufBC_2_D complexes for the wild-type and variant proteins as evidenced by SDS-PAGE, and circular dichroism spectra confirm the substitutions have not globally altered the protein structure (Figure S3). Persulfide transfer from SufE to SufBC_2_D can be measured by a persulfide trafficking assay in the presence of cysteine, SufS, and DTT (Figure 3A).(32) In this assay, persulfide formed on SufS is protected from reduction by DTT (solid line), the SufE persulfide intermediate is only partially protected (dashed line), and persulfide formed on SufBC_2_D is freely reduced by DTT and can be quantified using a methylene blue sulfide detection assay.

**Figure 2.**
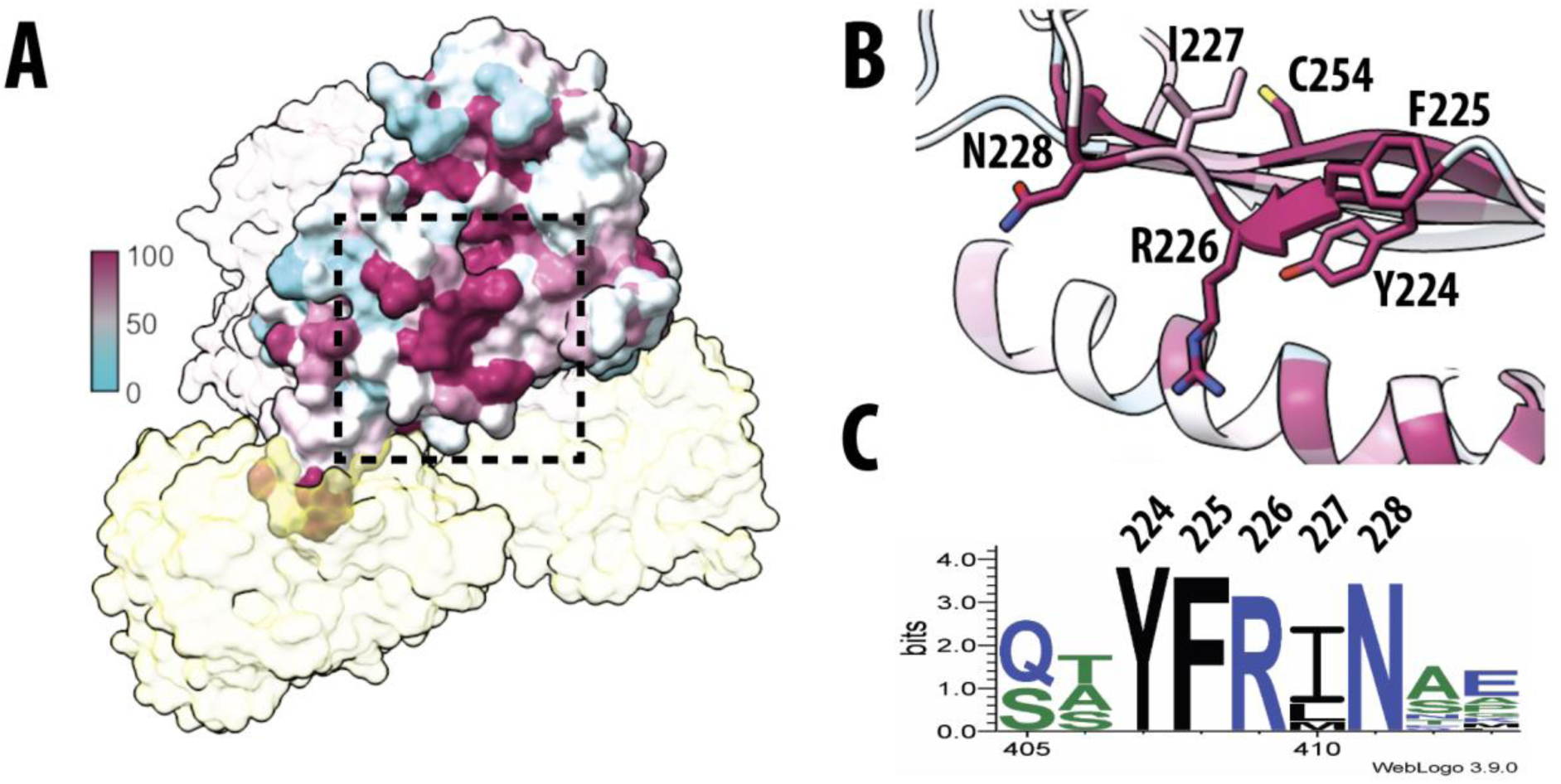
Visualization of SufB sequence conservation. (A) Percent sequence identity from the unbiased multiple sequence alignment visualized on the SufB subunit of SufBC_2_D (PDB ID: 5awf). The dotted box indicates the conserved patch under investigation. (B) Cartoon representation of the conserved patch with key residues labeled. Coloring is identical to panel A. (C) Hidden-Markov Model logo for the region shown in panel B with identical numbering.

**Figure 3.**
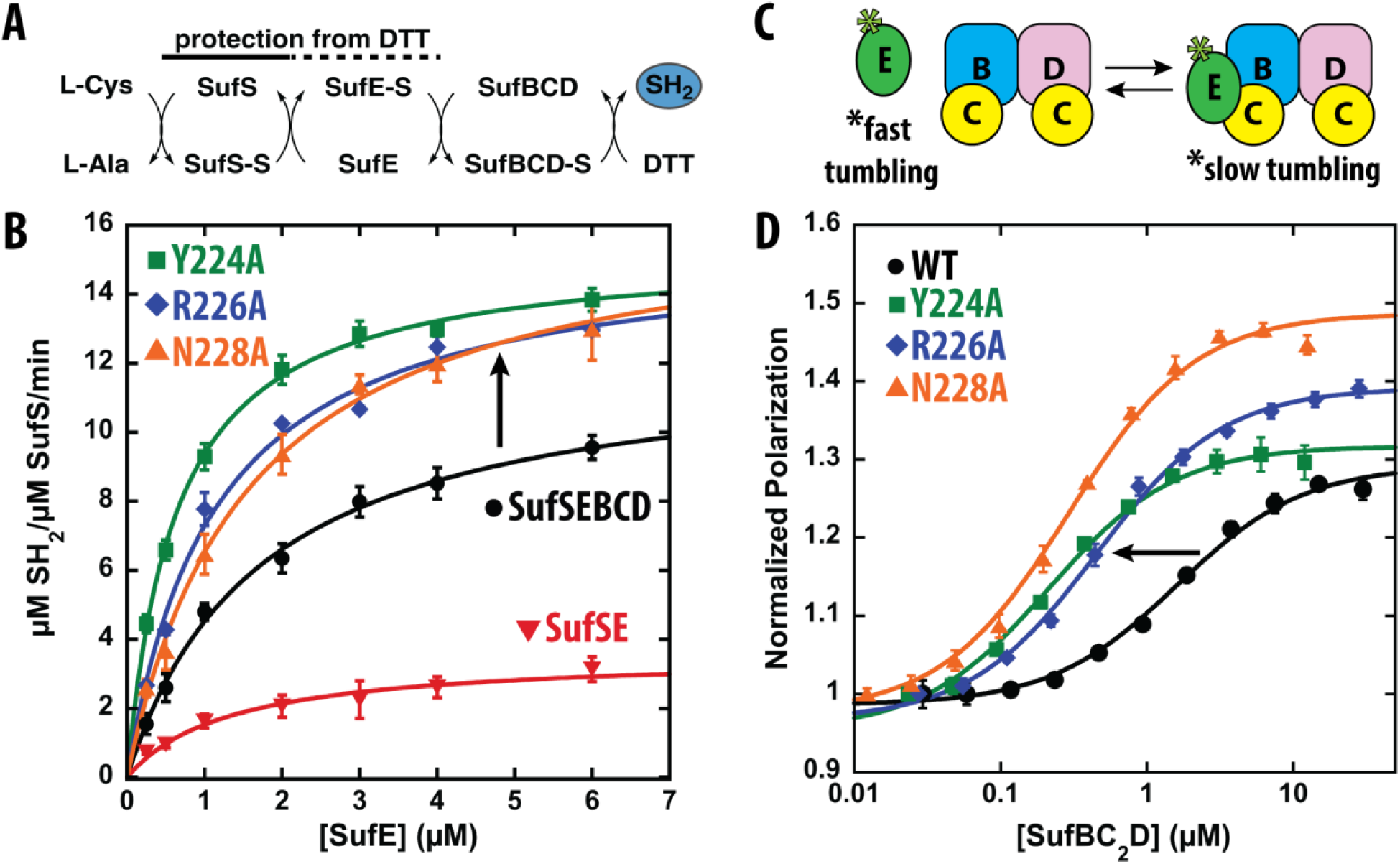
Functional characterization of SufB variants. (A) Schematic representation of the assay components and persulfide transfer pathway. The solid/dashed line represents the relative protection of each persulfide intermediate from reduction by DTT. Total sulfide released is detected using a methylene blue assay. (B) Rates of formation of sulfide determined at various concentrations of SufE (0-6 µM) and fixed concentrations of SufS and SufBC_2_D (0.25 µM). Kinetic parameters determined from the fits are shown in Table S1. (C) Schematic representation of the fluorescence polarization binding assay. C51A/E107C SufE was labeled with BODIPY-FL maleimide (*) and fluorescence polarization of the labeled SufE (0.05 µM) was titrated with SufBC_2_D (0-40 µM). Formation of the larger complex results in slower tumbling and increased polarization for the SufE fluorophore. (D) Fluorescence polarization binding data for wild-type SufBC_2_D and variants. The black arrow shows the shift in *K*_D_ values. Error bars are the results of standard deviation from triplicate data. Solid lines are from a fit of the data to Equation 2. *K*_D_ values determined from the fits are shown in Table S2.

In this assay, SufS and SufBC_2_D are held at equal concentrations (0.25 µM) and the concentration of SufE is varied to generate a pseudo-Michaelis-Menten curve. Varying SufE, rather than one of its two binding partners, prevents artificially shifting the equilibrium to either side of the reaction. Previously published control experiments showed that activation of SufS activity by SufBC_2_D requires both C51 on SufE and C254 on SufB.(28) With wild-type SufBC_2_D, the maximal rate of SufS activity is increased from 3.6 min^−1^ to 12.3 min^−1^ at saturating SufE concentrations (Figure 3B). Alanine substitutions at Y224, R226, and N228 result in ∼25-30% increases in maximal rates of persulfide formation (Figure 3B and Table S1). Previous characterization of R226A and N228A substitutions on SufB showed no increase in the rates of persulfide formation, likely due to differences in assay design, as those studies varied SufBC_2_D concentration, which may restrict SufE from freely sampling both binding partners.(29) Substitutions at F225 and I227, located between these residues, result in only modest increases in the rate of persulfide formation (Figure S4). Double variants at the 224 and 226 positions show that the effects are not additive (data not shown). The increased maximal rates of persulfide formation suggest that the substitutions stabilize a conformation in which the SufB C254 site is more accessible for transfer.

### Fluorescence polarization binding assays

We previously developed a C51A/E107C SufE variant that can be covalently labeled with BODIPY-FL-maleimide and used in fluorescence polarization binding assays (Figure 4C).(23) Using this technique, a *K*_D_ value of ∼2-3 µM was determined for SufE binding to wild-type SufBC_2_D.(28) Analysis of SufE binding to the SufBC_2_D variants shows that several variants (Y224A, R226A, and N228A) exhibit tighter binding to SufBC_2_D, with *K*_D_ values of 0.2-0.5 µM (Figure 4D and Table S2). The increase in binding affinity for SufE correlates with the increased rates of persulfide formation described above. The correlation between enhanced binding affinity and increased persulfide formation supports a model in which these substitutions stabilize a transfer-competent conformation of SufB that promotes productive interaction with SufE.

**Figure 4.**
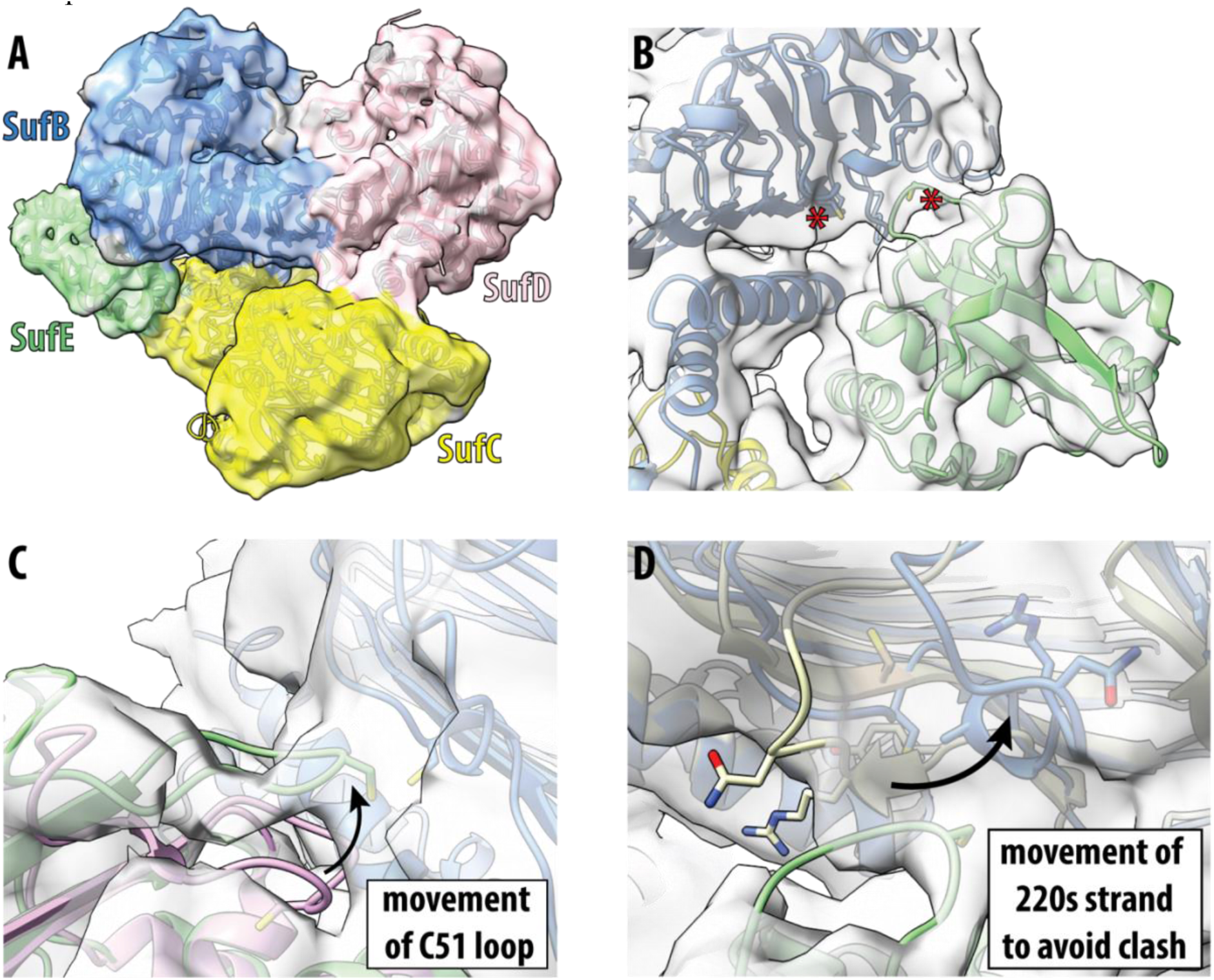
Cryo-EM structure of SufBC_2_D-SufE complex contoured at 4σ. (A) Map and model for the full complex with individual subunits labeled. (B) Detail view of SufB-SufE interactions. The two catalytic cysteine residues involved in persulfide transfer (SufE C51 and SufB C254) are rendered as sticks and highlighted with red asterisks. (C) Conformational changes in SufE are shown by superposition of the SufE monomer (purple, PDB ID: 1mzg) on the SufE subunit (green). Clear density is seen for the extended conformation of the C51 loop as opposed to the closed loop conformation of 1mzg. (D) Conformational changes in the SufB subunit (blue) shown by superposition with crystal structure of SufBC_2_D (tan, PDB ID: 5awf). Key residues on the SufB 220s loop are shown.

### SufB variants are not deficient in DTT-dependent Fe-S cluster assembly

The persulfide transfer and binding assays described above indicate that substitutions at positions 224, 226, or 228 do not disrupt functional interactions with SufE, ruling this out as an explanation for the reported in vivo requirement for these residues.(29) To determine whether the SufB variants impair in vitro Fe-S cluster assembly, anaerobic cluster assembly assays were performed with the wild-type and variant SufBC_2_D complexes. Equimolar amounts of SufS and SufE were mixed with a 100-fold excess of SufBC_2_D in the presence of ferrous ammonium sulfate and DTT in an anaerobic cuvette with a connected sidearm containing cysteine. Cluster assembly was initiated by rapidly mixing cysteine into the reaction mixture and monitored spectrophotometrically by following the increase at 420 nm, indicative of cluster formation. Wild-type SufBC_2_D reaches kinetic completion of cluster assembly after ∼4 hours, similar to previous reports (Figure S5). All three SufBC_2_D variants display nearly identical kinetics of cluster assembly under these conditions, despite their reported essential roles in vivo. Control reactions confirm that the C254 residue on SufB is required for cluster assembly (Figure S5). These results indicate that the effects of the SufB substitutions on persulfide transfer and SufE affinity are not due to global changes in scaffold function but instead reflect specific alterations in the persulfide transfer step.

### Cryogenic-electron microscopy captures the SufBC_2_D-E complex

The increase in binding affinity for SufE observed in the SufBC_2_D variants suggested they would be good candidates for structural characterization of the SufBC_2_D-E complex. Cryo-EM samples were prepared by mixing wild-type SufE with Y224A SufBC_2_D at a 5:1 molar ratio. Samples were blotted on cryo-EM grids and plunge-frozen. Single-particle images were collected from ∼4400 curated micrographs and analyzed in cryoSPARC (Figure S6). Initial blob picking and iterative 2D classification identified nine template classes, which were used to pick ∼1.6 million particles from the curated micrographs. Additional rounds of 2D classification resulted in a clean particle stack of 167,000 particles. Ab initio refinement identified three volumes, each of which were subjected to non-uniform refinement. Analysis of the three volumes identified compositional heterogeneity of the Suf proteins with a SufBC-D trimer (46k particles), a SufBC_2_D tetramer (43k particles), and a SufBC_2_D-E complex (76k particles). Subsequent analysis focused on the SufBC_2_D-E complex. Non-uniform refinement of the consensus SufBC_2_D-E particle stack resulted in a volume with an overall gold-standard Fourier shell correlation (FSC) resolution of 4.21 Å (Figure S7A). Although the overall resolution is moderate, the resulting map quality is comparable to the only previously reported X-ray structures of SufBC_2_D, which were refined at 4.28 Å and 2.96 Å.(26) Analysis of the map at a local resolution level shows high-resolution at the SufBC_2_D-SufE interface with lowest resolutions adjacent to the disordered region of SufB and distal regions of SufE (Figure S7B and S7C).

AlphaFold3 models independently predicted a SufBC_2_D-SufE arrangement similar to the experimentally observed complex(28), and were used as starting models for refinement in Phenix, followed by local remodeling in Coot to generate a final model (Figure 4A and Table S3). The cryo-EM map lacked density for residues 1-34 on the N-terminus and residues 94-152 of SufB. Similar regions are missing in the reported structures of the SufBC_2_D complex, suggesting that these regions are dynamic (Figure S1).(26) Several mechanistically relevant conformational changes are observed at the SufB-SufE interface. Starting with SufE, the most important change is a shift of the C51 loop (residues 48-54) to an extended conformation, relative to the inward position found in unbound SufE structures, such that the C51 residue is positioned toward SufB (Figure 4B). When the SufE structure (PDB ID: 1mzg) is superimposed on the SufE in complex, there is a clear region of map density for the extended loop and a lack of density in the position of the closed loop (Figure 4C). The shift in loop position is accompanied by an ∼8° splay of the adjacent β-strand 3 (residues 55-61) in the extended conformation relative to its position in the closed conformation.

In SufB, conformational changes in the 220s β-strand (residues 223-232), located exterior to the β-strand containing C254 (residues 247-257), allow accommodation of the extended C51 loop (Figure 4D). Notably, the orientation of the extended C51 loop places the persulfide transfer trajectory beneath the plane of the SufB-SufD axis rather than through the apparent opening generated by missing density in previous SufBC_2_D structures (Figure S1). This region partially adopts a helical conformation, repositioning the C-terminal portion of the strand to avoid a steric clash. A final conformational change is observed in the β-strand containing C254, which reorients from facing the interior of the β-sandwich to a more solvent-accessible position. Although neither cysteine is persulfidated, the sulfur atoms of C51 and C254 are positioned within ∼8 Å of one another suggesting only modest rearrangements would be required for persulfide transfer. Together, these conformational changes provide a structural basis for the enhanced binding affinity and persulfide transfer kinetics observed for the SufB variants and define how rearrangement of the 220s β-strand enables access to C254 for persulfide transfer. These observations support a model in which coordinated conformational changes in SufB and SufE enable productive, conformationally gated persulfide transfer to the SufB C254 site.

## Discussion

The molecular path of persulfide transfer in the *E. coli* Suf pathway from L-cysteine to the SufBC_2_D complex is broadly established (Figure 1B), with SufS cleaving the C-S bond of cysteine using a PLP cofactor to generate a covalent persulfide on C364. That persulfide is initially passed to C51 on SufE, followed by likely transfer to C254 on the SufB subunit of SufBC_2_D. These cysteine residues are evolutionarily conserved across SufE-dependent pathways in Gram-negative organisms as well as SufU-dependent pathways in Gram-positive organisms, suggesting a high degree of conservation for the sites of persulfide transfer. Despite the transient nature of the interactions, structures and mechanisms for the initial persulfide transfer by SufS are understood in both SufE-dependent and SufU-dependent systems. However, there is little mechanistic data concerning the transfer from either transpersulfurase to the SufBC_2_D scaffold. Here, we show that variants of SufB that stabilize a locally rearranged, transfer-competent conformation of SufBC_2_D enable structural characterization of the SufBC_2_D-SufE complex from *E. coli*, providing direct insight into this previously unresolved step.

### Conformational changes in the SufB subunit expose C254 for persulfide transfer

In previously reported structures for SufBC_2_D, the unresolved region of residues 78-158 created the appearance of an opening leading toward the C254-containing pocket that would require SufE to bind above the SufB-SufD axis (Figure S1).(26) In contrast, the current structure demonstrates that persulfide transfer from SufE occurs below the plane of the SufB-SufD interface axis and is enabled by localized conformational rearrangement of the 220s β-strand, which allows C254 to rotate out of the β-sandwich and into a solvent-accessible position compatible with transfer. Biochemical results with SufB variants suggest that residues Y224, R226, and N228 play a functional role in this conformational change as substitution at these positions altered SufB function whereas changes to the intervening residues F225 and I227 had little effect on function, arguing against a global disruption of the β-strand and instead supporting a localized structural role. The correlation between enhanced binding affinity and increased persulfide transfer rates suggests these substitutions stabilize a locally rearranged, transfer-competent conformation of SufB. Together, these findings resolve how SufE gains access to a previously buried cysteine and define a conformational mechanism for regulating access to the persulfide acceptor site.

### Complex formation drives conformational changes in SufE

In previous SufE structures, the C51 loop is found in a closed conformation with C51 facing the interior of the protein. This closed conformation likely explains the protected nature of the SufE persulfide intermediate but necessitates a conformational change to make the C51 residue available for persulfide transfer. We have previously identified R119 on SufE as a conserved molecular “check valve” responsible for sensing that persulfide transfer has occurred from SufS and participating in the dissociation of the SufS-SufE complex.(22) Subsequent studies on the SufS-SufU system from *B. subtilis* show that the analogous R125 interacts with the terminal sulfur of the BsSufU persulfide where it plays a similar mechanistic role in SufS-SufU complex dissociation.(24) In the current SufBC_2_D-SufE structure, the density suggests that R119 is oriented away from the C51 loop and towards the C-terminal helix of SufB where it can interact with several conserved acidic residues (E477 and E481). This repositioning is consistent with a model in which R119 participates in interfacial interactions that facilitate release of the C51 loop into an extended, transfer competent conformation. These observations suggest that R119 serves as a conserved regulatory element across multiple steps of the pathway, coupling conformational changes to directional transfer of the persulfide intermediate.

### Overall mechanism of persulfide transfer from SufE to SufBC_2_D

Based on the results presented here, we propose a refined mechanism for persulfide delivery from SufE to SufBC_2_D (Figure 5). SufE carrying persulfide can bind to SufBC_2_D (**1**) with a *K*_D_ value of ∼2-3 µM to form an initial interaction complex contacting SufB and SufC subunits (**2**), consistent with previous biochemical results.(27, 28) Conformational rearrangement of the 220s β-strand allow exposure of C254 and productive positioning of the SufE C51 loop, resulting in a higher affinity complex (*K*_D_ value ∼0.3 µM) (**3**). Concurrently, repositioning of R119 and associated interactions with SufB are proposed to enable extension of the C51 loop of SufE, placing the persulfide sulfur within transfer distance of C254. In the current structure, substitutions within the 220s β-strand appear to stabilize complex **3**. We hypothesize that this conformation represents a transient catalytic state normally populated during cluster assembly by intermediate steps, such as iron acquisition or flavin occupancy, that may precede sulfur transfer. Following transfer, ATP-dependent conformational changes involving SufC could disrupt the complex leading to loss of SufE (**4a**) consistent with the previously reported antagonistic feature of the two activities.(28) However, a simpler model where transfer of the persulfide to SufB is sufficient to promote dissociation of SufE (**4b**) cannot be ruled out. Following SufE dissociation, it is likely that the persulfide containing SufBC_2_D complex (**5**) exhibits unique conformational states to promote subsequent steps in cluster formation.

**Figure 5.**
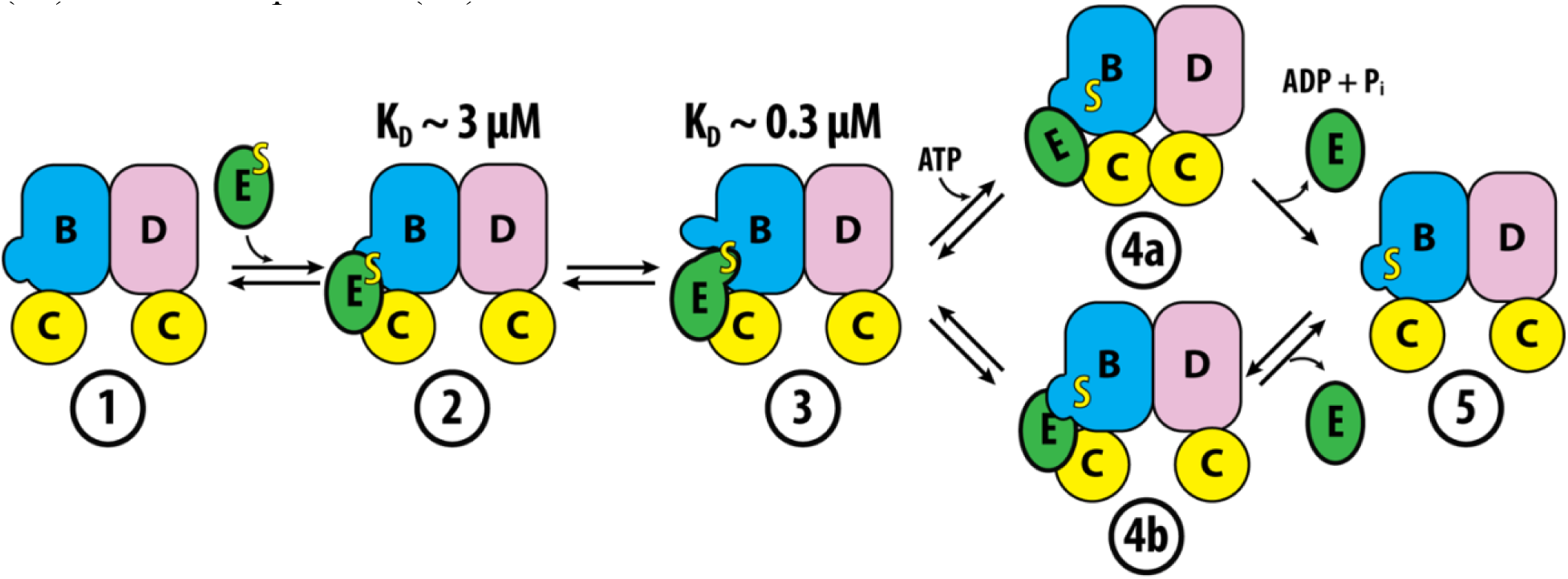
Proposed regulatory mechanism for persulfide transfer from SufE to SufBC_2_D. Persulfide-bound SufE initially forms a low-affinity complex with SufBC_2_D (**2**). An unidentified catalytic state promotes local conformational rearrangements that generate a transfer-competent state for persulfide transfer (**3**)n. Dissociation of SufE could occur through either an ATP-dependent (**4a**) or ATP-independent (**4b**) mechanism.

### A general mechanism for persulfide trafficking in the Suf pathway

A notable aspect of this model is that it provides a unified mechanistic framework for persulfide trafficking across the Suf pathway, in which a reactive covalent intermediate is passed between multiple proteins in the absence of a stable ternary complex or covalent tether. Fluorescence polarization assays show that SufE binds to both SufS and SufBC_2_D with similar affinity (∼2-5 µM), suggesting that partner selection is not governed solely by intrinsic binding preferences. Consistent with this model, SufE inhibits SufBC_2_D ATPase activity with a *K*_i_ value of ∼2 µM, and alkylation of the SufE active site cysteine does not substantially alter this affinity despite increasing affinity for SufS.(28, 33) These observations argue against a model in which directional trafficking is driven solely by preferential binding of persulfidated SufE to SufBC_2_D.

Structural comparison of SufE in complex with SufS and SufBC_2_D further suggests that persulfide transfer is mediated through a common catalytic face centered on the C51 loop. The newly identified SufB interfaces exhibit striking parallels to those previously characterized in the SufS-SufE complex (Figure 6A). For example, the α16 helix on SufS (residues 342-354) is important for forming the initial complex with SufE.(22) This helical interface structurally overlaps the N-terminal helix of SufB (residues 476-484) and bears a similar amino acid composition. For persulfide transfer from SufS to SufE, conformational changes in the flexible α3-α4 loop (residues 50-59) of SufS are responsible for creating the higher affinity state.(21) This flexible interface region occupies a structurally analogous position to the flexible 220s β-strand of SufB, which makes an analogous conformational change to promote persulfide transfer. In addition to the common interface, there are also distinct, non-overlapping surfaces on SufE used to engage each partner, with SufS interactions occurring at the C-terminal helix of SufE and interactions with the SufB-adjacent SufC subunit occurring through the N-terminal helix, consistent with previous biochemical data (Figure 6B).(28) This organizational architecture provides a structural basis for the inability of SufE to simultaneously bind SufS and SufBC_2_D, thereby preventing formation of a ternary complex and enforcing sequential transfer of the persulfide intermediate.

**Figure 6.**
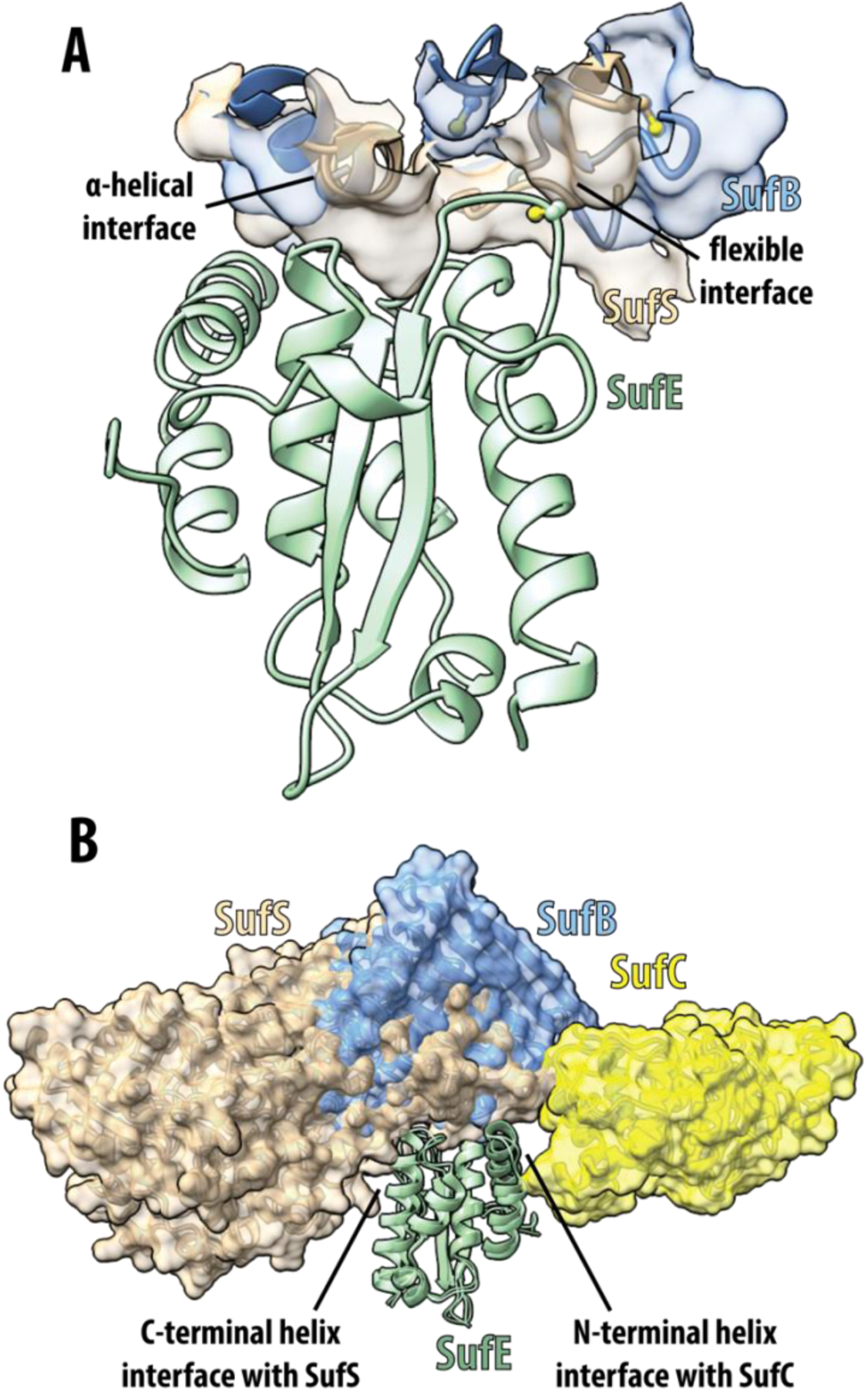
Structural parallels between SufS-SufE and SufBC_2_D-SufE complexes. (A) Superposition of SufE subunits (green, only one shown for clarity) from the SufS-SufE complex (PDB ID: 8vbs, tan) and the SufBC_2_D-SufE complex (blue) highlights analogous structural features involved in positioning the C51 loop for persulfide transfer. (B) Additional interfaces mediating SufS and SufBC_2_D binding are distinct and non-overlapping, preventing formation of a stable ternary complex.

Taken together, these findings support a general model in which conformational gating of transient protein-protein interactions governs persulfide transfer, ensuring both protection of reactive intermediates and directional sulfur flux through the pathway. While the specific catalytic intermediate that triggers this conformation of SufB remains to be identified, the ability to trap this state through amino acid substitutions highlights its functional relevance. A remaining question concerns the role of the unresolved region spanning residues 76-156 of SufB, which contains a CXXCXXXC motif shown to be non-essential in vivo.(29) If this region contributed directly to SufE binding, as predicted by AlphaFold3 models,(28) one might expect to observe density in the SufE-bound structure. The persistent absence of density suggests that this region remains highly flexible, even in the transfer-competent complex, and may play a role distinct from direct SufE engagement.

## Conclusions

Characterization of molecular mechanisms for bacterial Fe-S cluster assembly by the Suf pathway remains an important and understudied area, especially in comparison to the better understood Isc pathway. Here, the first structure of the SufBC_2_D-SufE complex defines how persulfide transfer to the scaffold is achieved through coordinated conformational changes in both SufB and SufE. These conformational changes expose the otherwise buried C254 acceptor site on SufB and promote extension of the SufE C51 loop, enabling productive transfer prior to cluster assembly. Integration of the structural and biochemical data supports a model in which persulfide transfer is regulated by conformational gating of both donor and acceptor proteins, providing a mechanism for controlling access to reactive sulfur intermediates. This gating mechanism offers a means to enforce directional sulfur transfer through a pathway that lacks a stable ternary complex, extending principles previously established for the SufS-SufE interaction to the SufE-SufBC_2_D step. Future studies will focus on how persulfide transfer from SufE integrates with subsequent SufBC_2_D-catalyzed steps in Fe-S cluster assembly, including ATP-dependent SufC dynamics, iron incorporation, flavin redox chemistry, and comparison of trafficking mechanisms in SufU-dependent Suf pathways.

## Experimental Procedures

### Sequence Similarity Network Generation

A representative sequence similarity network (34) was generated for the IPR010231 “Suf system FeS cluster assembly, SufB” domain from the InterPro database(35) using the Enzyme Function Initiative-Enzyme Similarity Tool (www.enzymefunction.org).(36) The ∼20,000 sequences were first reduced using UniRef90 families and further compacted using representative nodes at 80% sequence identity. The resulting network was downloaded as a Cytoscape readable .xgmml file for visualization.(37)

### Protein Growth and Expression

The previously described pDuet*SufBCD* vector(31) was used as template to generate the SufB variants C254A, Y224A, R226A, F225A, I227A, N228A, and the double variant Y224A/R226A via QuikChange PCR (Table S4). Following plasmid sequencing to confirm mutations and overall sequence integrity, the pDuet*SufBCD* plasmids were transformed into electrocompetent kanamycin resistant BL21(DE3)Δ*suf E. coli* cells for expression. In this cell line, the genomic *suf* operon has been replaced with a kanamycin resistance cassette to prevent contamination with endogenous wild-type proteins. The transformed cells were plated onto LB-agar plates with 100 µg/mL ampicillin and 50 µg/mL kanamycin and incubated overnight at 37 °C. A single colony from the plate was inoculated into 100 mL LB media with 100 µg/mL ampicillin and 50 µg/mL kanamycin and grown overnight at 37°C and 250 rpm. The next day, a 10 mL aliquot of the overnight culture was used to inoculate 1 L of sterile LB media containing 100 µg/mL ampicillin and 50 µg/mL kanamycin. Cells were grown at 37 °C and 250 rpm until OD_600_ reached 0.6, followed by induction with 0.5 mM isopropyl-β-D-thiogalactoside (IPTG). After growth for 3 hours post-induction, cells were harvested by centrifugation for 10 minutes at 10,000 x g. Harvested cells were stored in −80 °C until purification. SufS and SufE were expressed as previously described.(23)

### SufBC_2_D Purification

Frozen cell pellets were resuspended in lysis buffer (50 mM Tris, pH 7.5, 500 mM NaCl, 20 mM imidazole, 10 mM β-mercaptoethanol (BME), 1 mM phenylmethylsulfonyl fluoride (PMSF), 1 mM MgCl_2_, and 2 µg/mL DNase. The resuspended cells were lysed via sonication for 10 minutes with 30 seconds cycles of on/off, and insoluble debris was removed by centrifugation. The lysate solution was then loaded onto the 5 mL HisTrap column (Cytiva) and washed with buffer A (50 mM Tris HCl pH 7.5, 500 mM NaCl, 20 mM imidazole, and 10 mM BME). A linear elution gradient with buffer B (50 mM Tris HCl pH 7.5, 500 mM NaCl, 500 mM imidazole, and 10 mM BME) included a five-column-volume hold at 10% B, during which SufBC_2_D eluted. Fractions containing SufBC_2_D were concentrated and loaded onto Superdex 200 column for further purification with running buffer of 25 mM 3-(N-morpholino)-propanesulfonic acid (MOPS) pH 7.5, and 150 mM NaCl. SufBC_2_D elutes as a single peak and protein purity was assessed by SDS-PAGE. Fractions were pooled, concentrated, and stored in 10 % glycerol in −80 °C for further characterization. The concentration of SufBC_2_D was measured using the calculated extinction coefficient (ε_280_ = 130,070 M^−1^cm^−1^). Purified SufS and SufE proteins required for kinetic and binding assays described below were purified and stored as previously described.(22)

### Persulfide trafficking assay

To determine if SufB variants can activate SufSE cysteine desulfurase activity to the same extent as WT SufBC_2_D, a methylene blue assay was performed to quantify the rates of sulfide formation by SufS in the presence of SufE and SufBC_2_D.(29, 32) The assay was carried out at 25 °C and contained 50 mM MOPS pH 7.5, 150 mM NaCl, 0.25 μM SufS, 0.25 μM SufBC_2_D or its variants, 2 mM cysteine, varying amounts of SufE (0 – 6 μM), and 2 mM DTT. The reaction was initiated by addition of cysteine, and 200 μL aliquots were quenched at various time points with 25 μL of 25 mM N,N-dimethyl-p-phenylenediamine (DMPD) in 7.2 M HCl followed by addition of 2 μL 30 mM FeCl_3_ in 1.2 M HCl. The labeled mix was incubated for 30 minutes at room temperature and transferred to a clear 96-well plate. Absorbance at 670 nm was measured on BioTek Synergy2 multi-well plate reader. A sodium sulfide standard was made under similar conditions and used to calculate the amount of sulfide generated in the enzyme reaction. Linear progress curves for sulfide production were determined from at least four time points. Kinetic parameters were determined from plots of initial velocity versus [SufE] fit to the Michaelis-Menten equation.

### Fluorescence polarization SufE binding assay

The fluorescently labeled SufE C51A/E107C (SufE DM) was prepared as described previously.(23) Labeled SufE DM (50 nM) was titrated against varying concentrations of SufBC_2_D and its variants (40 μM – 0.01 μM) in a buffer containing 50 mM MOPS, 150 mM NaCl, pH 8 and 0.1 mg/mL bovine serum albumin. The samples were incubated in the dark for 30 minutes. After incubation, the fluorescence polarization (480 nm excitation, 520 nm emission) was measured using a BioTek Synergy2 multiwell plate reader. After plotting fluorescence polarization versus [SufBC_2_D], *K*_D_ values were determined by fits to the quadratic binding equation.(38)

### Iron-sulfur cluster reconstitution

The iron-sulfur cluster assembly reactions were performed anaerobically and monitored on a photodiode array spectrophotometer for around 6 hours for WT SufBC_2_D and its variants. Reactions conditions were 50 mM MOPS, 150 mM NaCl pH 7.5 containing 0.5 μM SufS, 0.5 μM SufE, 20 μM SufBC_2_D, 500 μM cysteine, 100 μM Fe(NH_4_)_2_(SO_4_)_2_•6H_2_O, and 2 mM DTT. The reaction mixture was assembled in an anaerobic cuvette with a separate bulb containing cysteine and made anaerobic on a Schlenk line with alternating cycles of vacuum and argon, then incubated at 25 °C for 10 minutes. Cluster assembly was initiated by mixing cysteine into the cuvette contents and monitored on photodiode array spectrophotometer over the 6-hour time course.

### Structure determination of SufBC_2_D-SufE complex by cryo-EM

Holey carbon grids with copper support were glow discharged on a Pelco EasiGlow. Purified SufE (12.5 µM) was combined with SufBC_2_D (2.5 µM) containing the Y224A substitution on SufB subunits (Y224A SufBC_2_D). Grids were blotted and plunge frozen in liquid ethane using a Vitrobot Mark IV and cryo-EM movies were collected using a Glacios 2 with a Falcon 4i direct detector. The movies were processed and single particle reconstruction was performed using CryoSPARC v4 (Figure S6).(39) Non-uniform refinement of a final SufBC_2_D-SufE particle stack resulted in a 4.21 Å reconstruction as determined by gold-standard Fourier shell correlation plots. An initial molecular model was generated using AlphaFold3(40) and docked using ChimeraX(41) followed by rigid-body refinement in Phenix(42) and loop remodeling in Coot(43) to arrive at a final molecular model.

## Data Availability

All kinetic and biophysical data are contained within the manuscript and supporting information. Cryo-EM volume data have been deposited in the EMDB under accession code EMD-75839. Structural data have been deposited in the PDB under accession code 11MP.

## Supporting Information

This article contains supporting information

## Acknowledgements

We thank Dr. James Kizziah (University of Alabama at Birmingham Cryo-EM Facility) for his assistance with cryo-EM sample preparation and data collection.

## Author Contributions

N.C. and N.M. contributed to investigation. N.C., J.A.D., and P.A.F. contributed to formal analysis and writing. J.A.D. and P.A.F. contributed to supervision.

## Abbreviations

AF3: AlphaFold3 Cys, cysteine
DTT: dithiothreitol
EFI-EST: Enzyme Function Initiative-Enzyme Similarity Tool
Fe-S: iron-sulfur
FSC: Fourier shell correlation
HMM: Hidden-Markov Model
*K*_D_: equilibrium dissociation constant
SSN: sequence similarity network

