## Supporting Information for "The structure of the SufBC_2_D-SufE complex reveals the mechanism of sulfur transfer in bacterial Fe-S cluster assembly"

**Table S1. Fit parameters for persulfide trafficking assay data.** Initial velocities were determined in triplicate. Kinetic parameters were determined from a fit to the Michaelis-Menten equation. Data are shown in Figure 3B of the main manuscript.

| reaction conditions | $k_{cat}$ ($min^{-1}$) | $K_{0.5}$ |
|---|---|---|
| SufS-SufE alone | 3.6 ± 0.3 | 1.3 ± 0.3 |
| $SufBC_2D$-SufS-SufE | 12.3 ± 0.4 | 1.7 ± 0.2 |
| Y224A $SufBC_2D$-SufS-SufE | 15.3 ± 0.3 | 0.63 ± 0.04 |
| F225A $SufBC_2D$-SufS-SufE | 11.7 ± 0.5 | 1.0 ± 0.1 |
| R226A $SufBC_2D$-SufS-SufE | 15.6 ± 0.7 | 1.1 ± 0.1 |
| I227A $SufBC_2D$-SufS-SufE | 14.4 ± 0.8 | 1.6 ± 0.2 |
| N228A $SufBC_2D$-SufS-SufE | 17.3 ± 0.7 | 1.7 ± 0.2 |

**Table S2. Fit parameters for SufE fluorescence polarization binding assays.** Binding curves were generated in triplicate. Dissociation constants were determined by fits of the data to the quadratic binding equation. Data are shown in Figure 3C of the main manuscript.

| SufB variant | $K_d$ (μM) |
|---|---|
| WT | 1.3 ± 0.2 |
| Y224A | 0.15 ± 0.02 |
| F225A | 1.5 ± 0.1 |
| R226A | 0.45 ± 0.05 |
| I227A | 1.8 ± 0.2 |
| N228A | 0.29 ± 0.02 |

**Table S3. Cryo-EM data collection, refinement, and validation statistics**

| **Data Collection** | **$SufBC_2D$-SufE** |
|---|---|
| PDB ID | 11MP |
| EMBD ID | EMDB-75839 |
| microscope | Thermo Fisher Glacios |
| detector | Falcon 4i |
| magnification | 190,000 |
| voltage (kV) | 200 |
| exposure ($e^-/Å^2$) | 60 |
| defocus range (μM) | -0.01 to -3.3 |
| raw pixel size (Å) | 0.73 |
| number of movies | 5500 |
| initial particles | 1.58 M |
| imposed symmetry | $C_1$ |
| final particle number | 76,000 |
| resolution (Å) | 4.21 |
| FSC threshold | 0.143 |
| **Refinement** | |
| map pixel size (Å) | 0.73 |
| model resolution (Å) | 4.39 |
| FSC threshold | 0.143 |
| *B* factors ($Å^2$) | |
| protein | 313.39 |
| r.m.s. deviations | |
| bond lengths (Å) | 0.12 |
| bond angles (°) | 0.32 |
| validation | |
| MolProbity score | 1.89 |
| clash score | 15 |
| rotamer outliers (%) | 0.0 |
| Ramachandran plot | |
| favored (%) | 97 |
| allowed (%) | 3 |
| disallowed (%) | 0 |

**Table S4. Mutagenesis primer list**

SufB variants generated using WT pDuet*SufBCD* plasmid.

| Variant | | Primer (5' - 3') |
|---|---|---|
| Y224A | Forward | ttctgcgttaatgcgaaaagcggtggaaagttccatcggg |
| Y224A | Reverse | cccgatggaactttccaccgcttttcgcattaacgcagaa |
| F225A | Forward | tgcgttaatgcgagcataggtggaaagttccatcgggcag |
| F225A | Reverse | ctgcccgatggaactttccacctatgctcgcattaacgca |
| R226A | Forward | cggtttttctgcgttaatggcaaaataggtggaaagttccatc |
| R226A | Reverse | gatggaactttccacctattttgccattaacgcagaaaaaaccg |
| I227A | Forward | gcccggtttttctgcgttagcgcgaaaataggtggaaagtt |
| I227A | Reverse | aactttccacctattttcgcgctaacgcagaaaaaaccgggc |
| N228A | Forward | gatggaactttccacctattttcgcattgccgcagaaaaaaccgg |
| N228A | Reverse | ccggtttttctgcggcaatgcgaaaataggtggaaagttccatc |

SufB variants generated using R226A pDuet*SufBCD* plasmid.

| Variant | | Primer (5' - 3') |
|---|---|---|
| R226A/Y224A | Forward | ttctgcgttaatggcaaaagcggtggaaagttccatcggg |
| R226A/Y224A | Reverse | cccgatggaactttccaccgcttttgccattaacgcagaa |

**Figure S1. Potential access to C254 on SufB.** Analysis of $SufBC_2D$ (PDB ID: 5awf) shows potential access to C254 for persulfide transfer. Unresolved density for SufB residues 80-156 (magenta line) suggest that this access point (dashed circle) is blocked in the complete structure. According to the model, SufE would deliver persulfide from a position above the SufB-SufD axis.

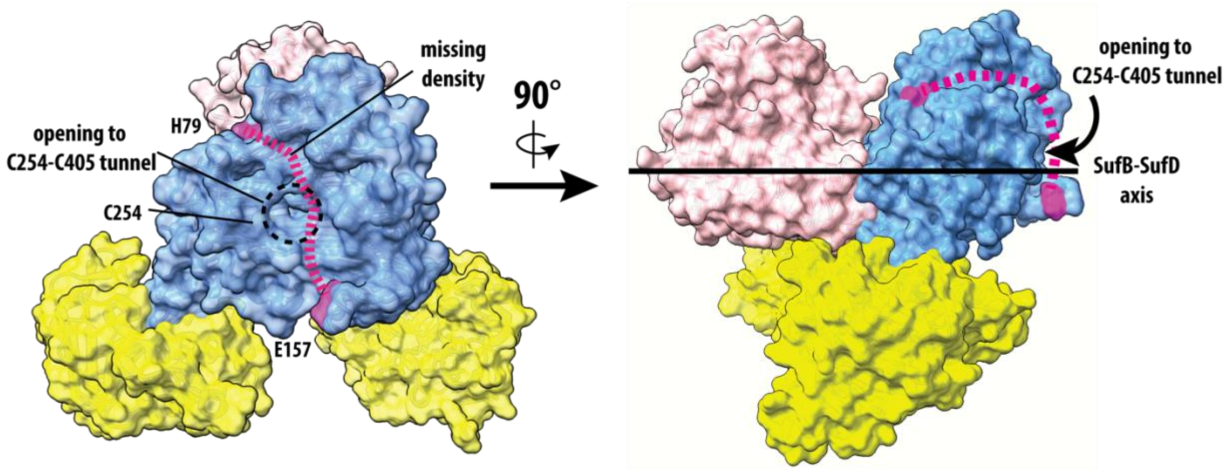

**Figure S2. Sequence similarity network and multiple sequence alignment for SufB.** (A) Sequence similarity network generated from representative bacterial SufB sequences in IPR010231. Nodes represent individual protein sequences and edges represent pairwise similarity relationships above the selected threshold. (B) Multiple sequence alignment highlighting conservation. Conserved residues are shaded according to sequence identity. The 220s strand is highlighted with a red box, and C254 is denoted with a red circle.

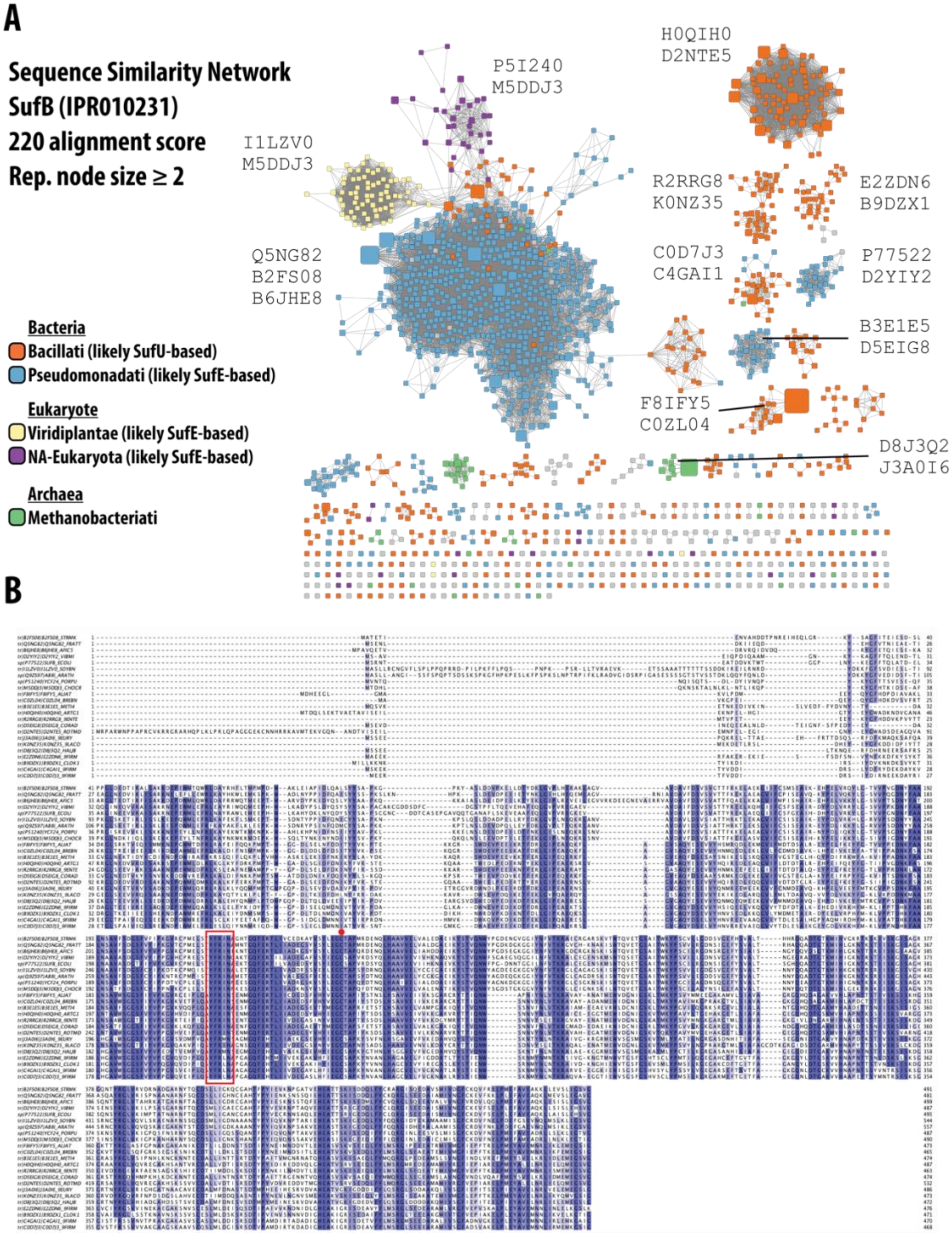

**Figure S3. Quality controls for final wildtype and variant SufBC$_2$D proteins.** (A) 12% SDS-PAGE gel showing purity and subunit composition. (B) Circular dichroism spectra of WT SufBC$_2$D and its variants were acquired in the far-UV region (190-280 nm) at 25°C using a spectropolarimeter (J-1500 CD Spectrometer). Each spectrum is an average of five scans and was baseline corrected using a 50 mM phosphate buffer pH 8.0 as reference. The spectral data was normalized by area to eliminate the changes in each spectrum due to changes in protein concentrations and enable an effective way of comparison of the protein shapes. For the final concentration, protein samples were diluted to 0.2 mg/ml using 50 mM phosphate buffer pH 8.0 to minimize the interference of 150 mM NaCl with the CD spectra in the far-UV region (190-280 nm).

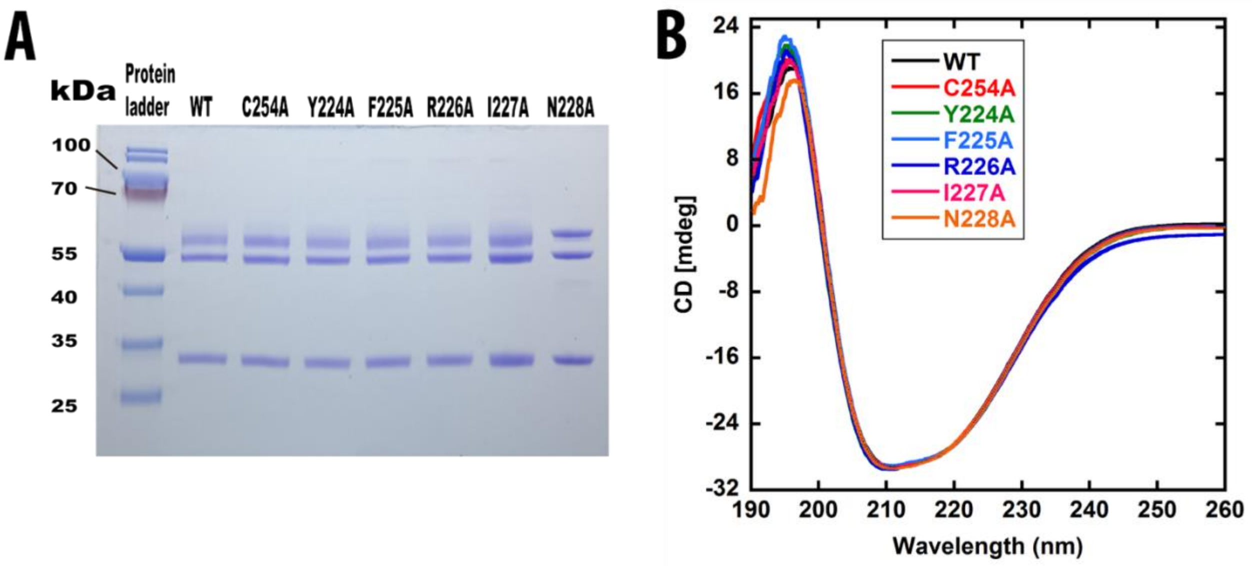

**Figure S4. Biochemical results for additional SufBC$_2$D variants F225A and I227A.** (A) methylene blue assay for persulfide transfer (B) fluorescence polarization results.

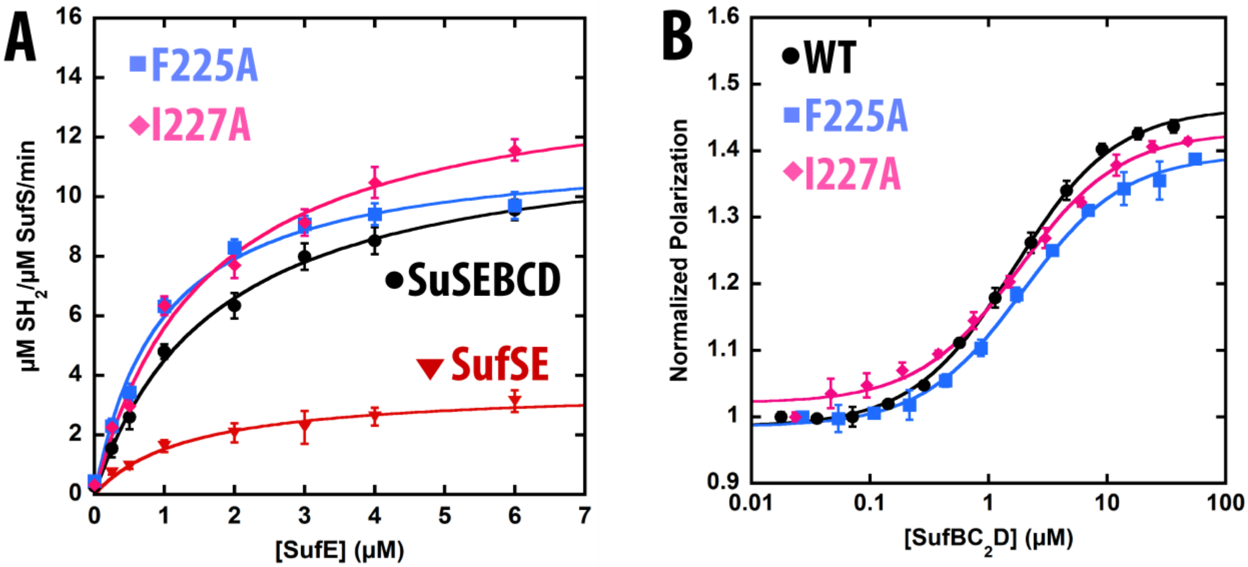

**Figure S5. In vitro Fe-S cluster assembly assays.** Cluster assembly was monitored using a photodiode array spectrophotometer. Assays for each protein were assembled as described in the methods section. Increases in absorbance at 420 and 620 nm are consistent with formation of a [4Fe-4S] cluster as previously characterized. Plots are labeled based by substitutions in the SufB subunit of the $SufBC_2D$ complex, with the C254A substitution used as a negative control.

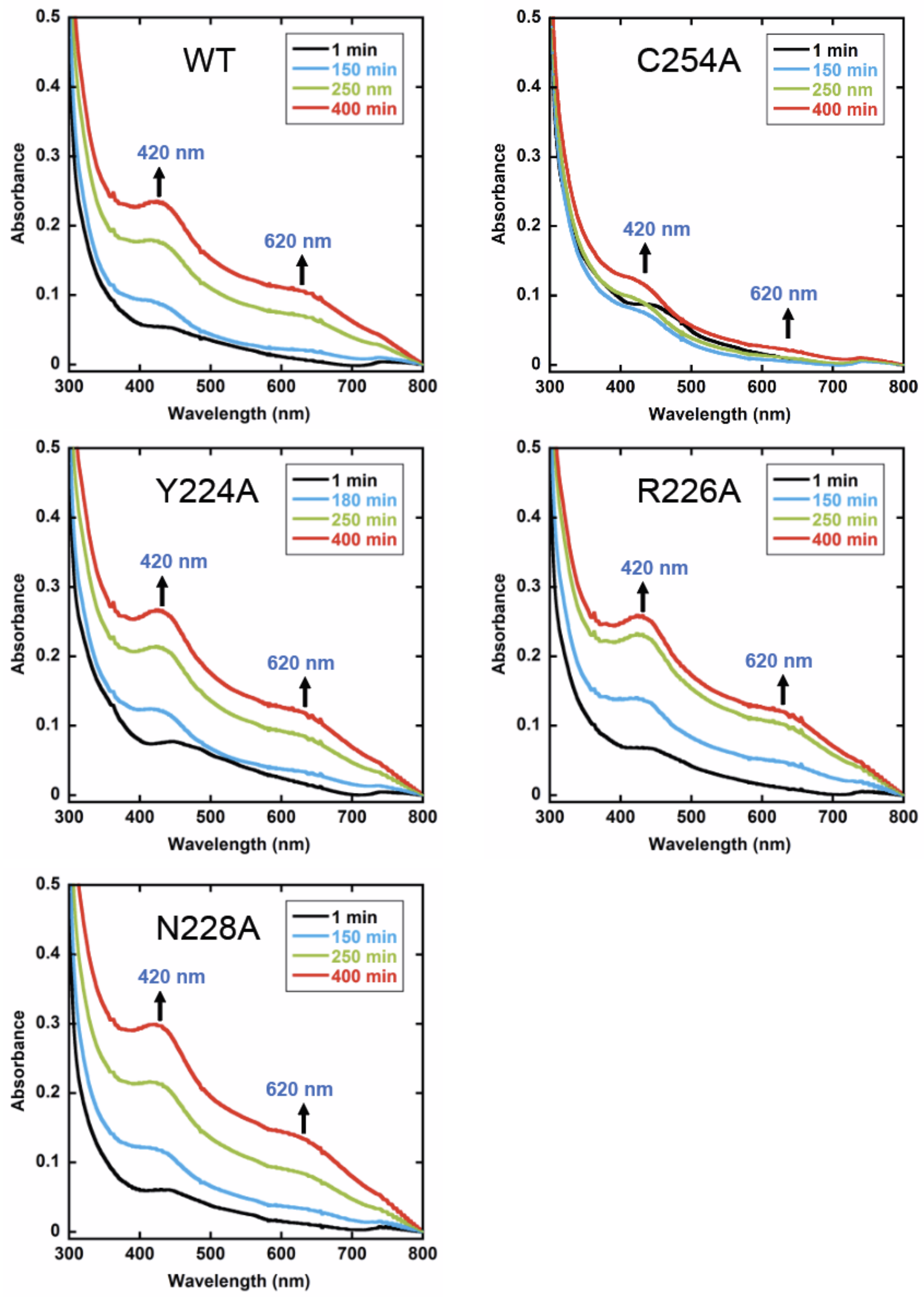

**Figure S6. Cryo-EM processing workflow.** All steps were performed in CryoSPARC version 4.0.

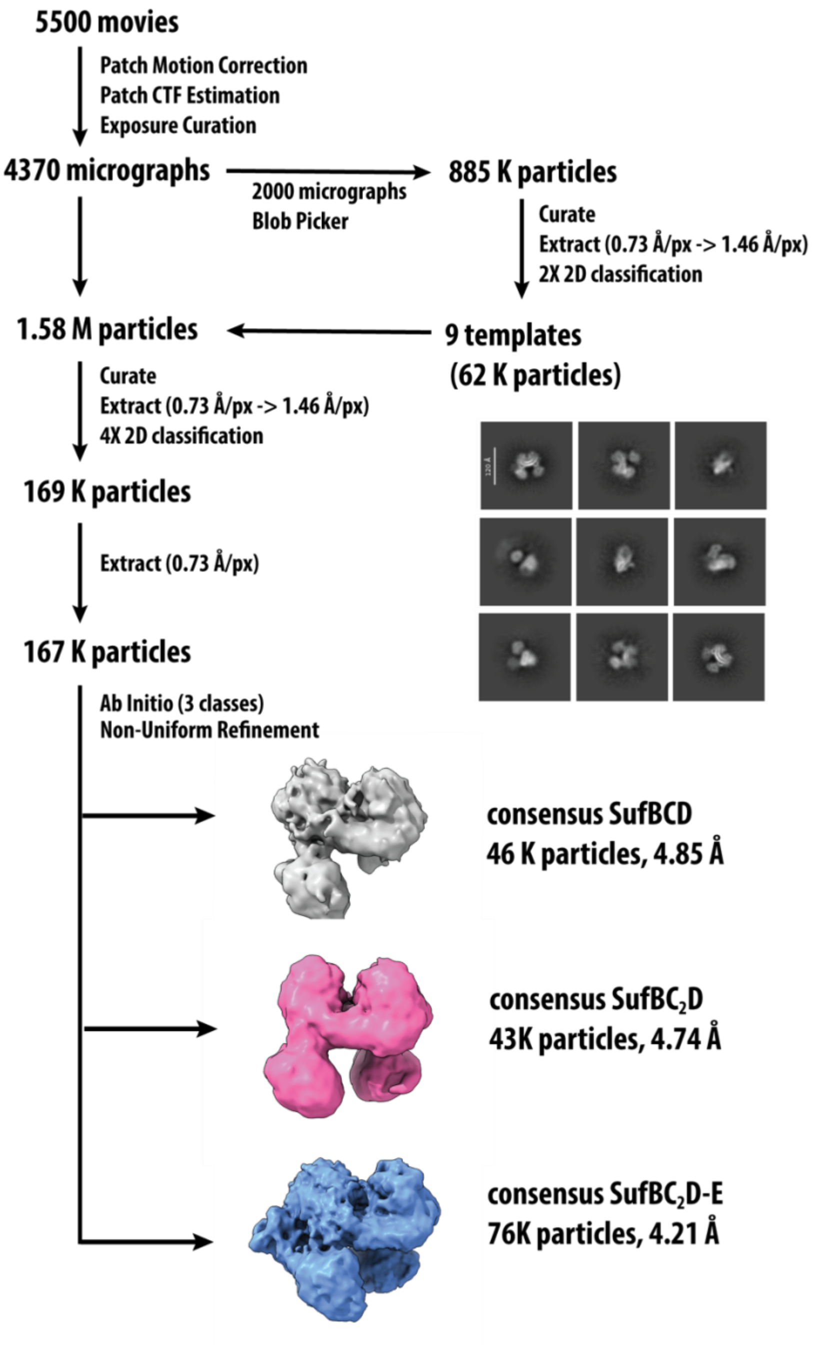

**Figure S7. Resolution analysis of the $SufBC_2D$-SufE cryo-EM reconstruction.** (A) Gold-standard Fourier shell correlation (FSC) curve used to estimate overall map resolution at the 0.143 criterion. (B) Local resolution map colored according to estimated local resolution values, demonstrating variation in map quality across the complex. Regions corresponding to flexible peripheral domains exhibit lower local resolution relative to the core scaffold architecture. (C) Detail view of SufB-SufE interface outlined in dashed box from panel B.

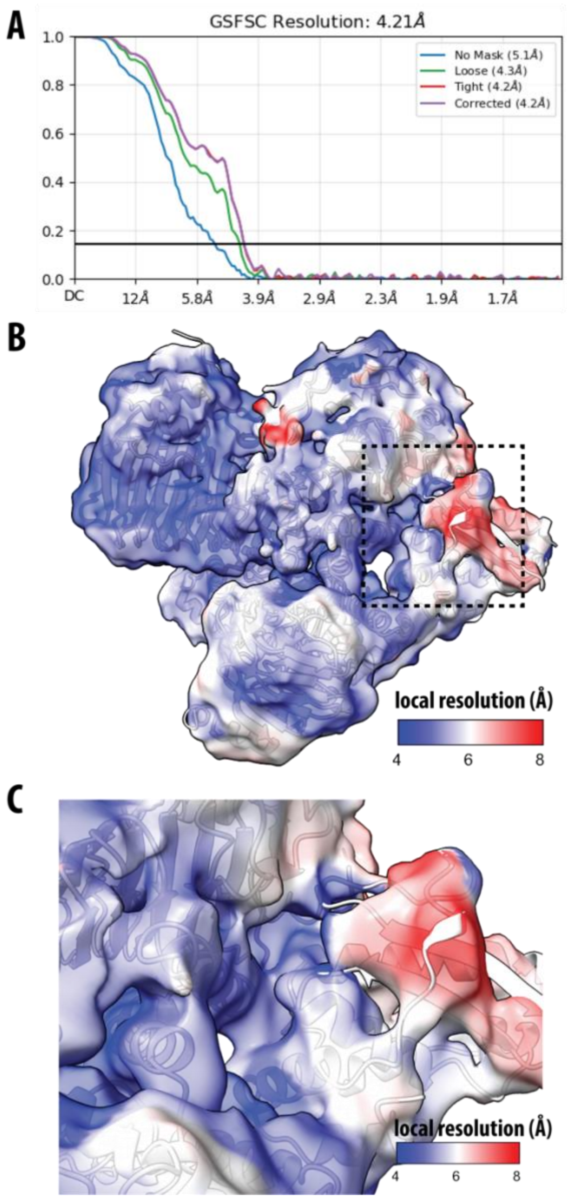
